# Intricate structure of the interphase chromocenter revealed by the analysis of a factor involved in species formation

**DOI:** 10.1101/441675

**Authors:** Natalia Y. Kochanova, Tamas Schauer, Grusha Primal Mathias, Andrea Lukacs, Andreas Schmidt, Andrew Flatley, Aloys Schepers, Andreas W. Thomae, Axel Imhof

## Abstract

In higher eukaryotes centromeres often coalesce into a large intranuclear domain called the chromocenter. Chromocenters are important for the organization of pericentric heterochromatin and a disturbance of their formation results in an upregulation of repetitive elements and causes defects in chromosome segregation. Mutations in the gene encoding for the centromere associated *Drosophila* speciation factor HMR show very similar phenotypes suggesting a role of HMR in chromocenter architecture and function. We performed confocal and super resolution microscopy as well as proximity based biotinylation experiments of HMR and its associated protein HP1a to generate a molecular map of HMR and HP1a bound chromatin. Our work reveals an intricate internal structure of the centromeric chromatin region, which suggests a role of HMR in separating heterochromatin from centromeric chromatin.

## INTRODUCTION

In many eukaryotes pericentromeric and centromeric chromatin of multiple chromosomes cluster in interphase to form a nuclear domain, the so-called chromocenter (1-3). The chromocenters are composed of pericentromeric heterochromatin and CenpA containing centromeric chromatin, which are both rich in repetitive DNA and evolutionarily highly dynamic (4, 5). They form and are held together by multiple components ranging from RNA transcribed from the repeats (6) over DNA binding factors (7), protein-protein interactions (8) to histone posttranslational modifications (9). Interference with chromocenter formation results in an upregulation of transposable elements, mitotic defects and the formation of micronuclei (7, 8). Despite their functional importance, the centromere as well as the pericentromeric repeats are highly divergent with regard to size, sequence and protein composition even in very closely related species (4, 10, 11). This rapid divergence of centromeric sequences is thought to be accompanied by an adaptive evolution of centromere binding proteins to counteract a meiotic drive, which would otherwise result in the potentially deleterious expansion of centromeric repeats (5, 12). This hypothesis is supported by the fact that many of the gene products that result in hybrid incompatibility are either proteins that bind to the centromeric or pericentromeric heterochromatin (13-16) or RNA molecules that localize in this region (17, 18). One of the best characterised hybrid incompatibility factors is the Hybrid male rescue protein HMR (19). HMR interacts with the heterochromatin protein HP1a and co-localizes with the centromere-specific H3 variant dCenpA (CID) in *Drosophila* cell lines and imaginal disc cells (13). HMR binding is also observed at several euchromatic sites where it colocalizes with known boundary factors (20). Interestingly, the intracellular localization of HMR varies among different tissues. In interphase cells of larval brains it is primarily found at pericentromeric heterochromatin (15, 21, 22) whereas it also associates with telomere structures on salivary gland polytene chromosomes (13, 22). Mutations of *Hmr* results in an upregulation of transposable elements (13, 23) and an increase in mitotic defects (13). As this phenotype is very similar to defects upon knockdown of nucleoplasmin (Nlp), which interacts with HMR and plays a role in centromere clustering (8, 13), we wondered about HMR’s role in chromocenter architecture. To investigate the intricate structure of the *Drosophila* chromocenters we performed confocal and super resolution microscopy using antibodies directed against HP1a, HMR and dCenpA and determined the proteome in proximity of HMR and HP1a using APEX2 based proximity biotinylation. Our results suggest that HMR is located at boundaries between HP1a containing heterochromatin and centromeric or transcriptionally active chromatin. Besides the proximity to heterochromatic and known centromeric factors we also observe a close proximity of HMR to nucleolar proteins, transcription factors, nuclear pore components and the condensin and cohesin complex. These findings suggest an important role of HMR in orchestrating the formation of the evolutionarily very dynamic chromocenter domain. As a consequence, the differential regulation of HMR in different species of *Drosophila* results in complex mitotic defects in hybrid animals containing two different and separately evolved genomes.

## RESULTS

### HMR and dCenpA form an interdigitated centromeric network

To confirm the previously detected centromeric localization of HMR in *Drosophila* cells with another antibody, we performed immunofluorescent staining using a FLAG antibody in a cell line where HMR is endogenously tagged with the FLAG epitope at the C-terminus using CRISPR/Cas9 (20). Consistent with our previous results (13) most of the FLAG signals co-localize with the signal we obtained when using an anti-dCenpA antibody, which confirms the close proximity of HMR and centromeric chromatin during interphase (Figure 1A). Notably, the quantitation of more than 120 interphase centromeres indicated that 57% of all centromere foci overlap with HMR foci. This suggests that the localization of HMR to the centromere is either cell cycle regulated or specific for a subset of centromeres (Figure 1B), which may explain the differences observed when staining HMR in different tissues (13, 15). Another observation that is difficult to reconcile with the clustered staining of HMR in cells is the fact that ChIP-seq data suggest HMR binding along the euchromatic arms often at boundaries between HP1a containing heterochromatin and actively transcribed genes (20). As centromeres are also transcribed (10) and in proximity to pericentromeric heterochromatin, we wondered if HMR might also localize to a putative boundary between these two different chromatin regions. Due to its repetitive nature, pericentromeric as well as centromeric chromatin is difficult to map in ChIP-seq experiments. We therefore investigated the interphase centromere and its surrounding chromatin using super resolution STED microscopy. These experiments revealed that the chromocenter is not homogenously formed by dCenpA containing chromatin but constitutes a structural meshwork of interdigitated dCenpA and HMR proteins (Fig. 1C, Movie S1), which we assume to constitute an intranuclear domain frequently defined as the chromocenter (24). In agreement to previous results (25), this centromeric region is surrounded by large lobes of HP1a containing heterochromatin, which are often bordered by HMR signals (Supplemental Fig. S1, Movie S1). This intricate structure is further confirmed by super resolution microscopy stainings of dCenpA and dCenpC as well as HMR and dCenpC. These co-stainings revealed a substantial colocalization of dCenpC with dCenpA, in contrast to the shifted localization observed between HMR and dCenpC or dCenpA (Fig. 1D).

**Figure 1:**
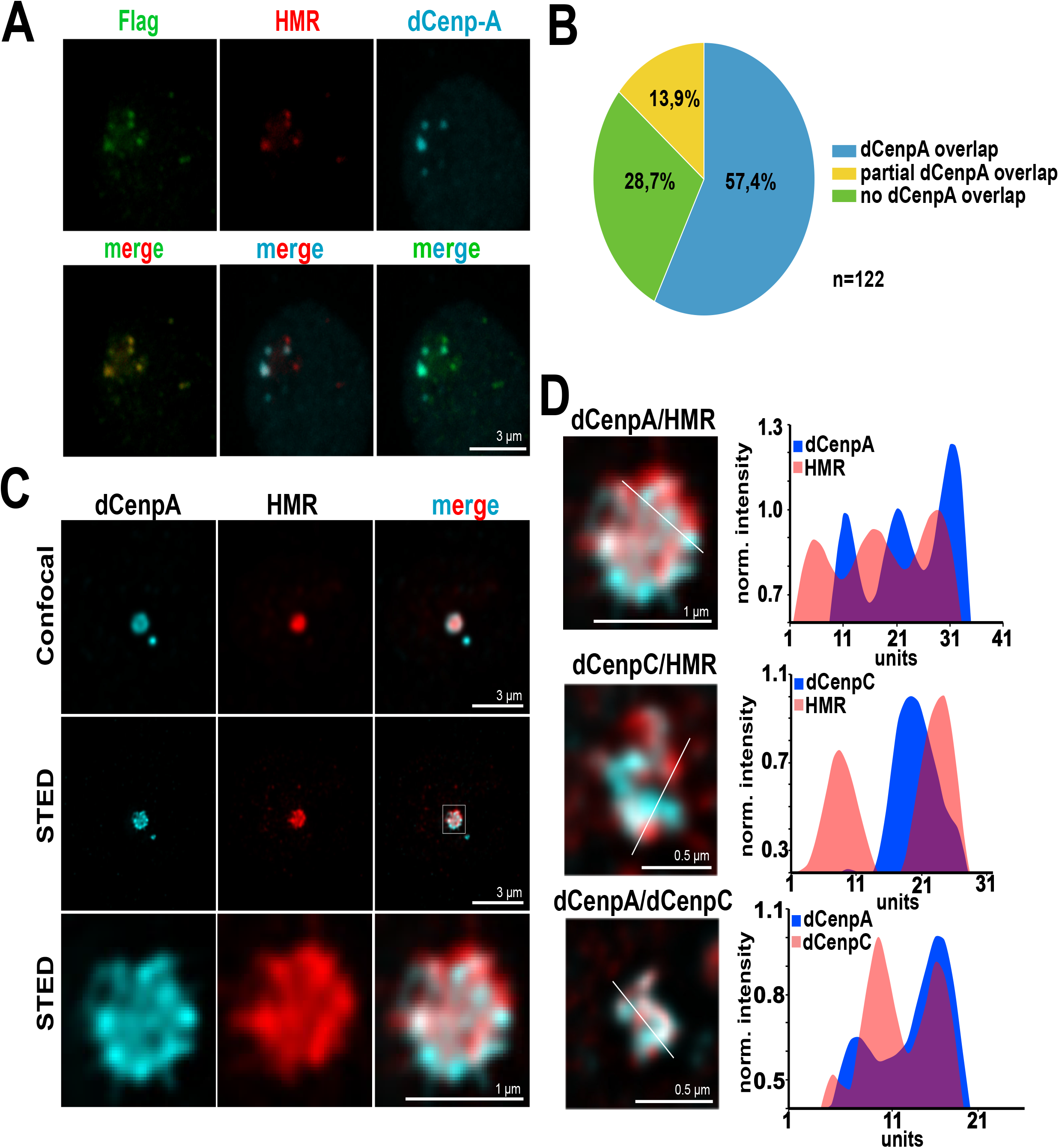
(**A**) – Staining of S2 FLAG-HMR line (20) with mouse anti-FLAG (Sigma), anti-HMR 2C10 and rabbit anti-dCenpA antibodies. (**B**) Quantification of centromeres overlapping with HMR. Experiments were performed in a dCenpA-GFP expressing cell line (53). In total 25 cells and 122 centromeres identified using an anti GFP antibody were counted. (**C**) Confocal (top panel) and stimulated emission depletion (STED) microscopy (middle panel) images of dCenpA and HMR. The bottom panel shows a zoom in of the centromeric region (**D**) Plot profiles of the highest fluorescence intensities of STED images of the chromocenter from S2 cells using anti dCenpA, anti dCenpC and anti HMR antibodies. Intensity graphs were built in ImageJ and normalized to one of the maximum peaks.

### HMR and HP1a APEX2 fusion proteins localize similarly to the endogenous proteins

To further unravel the details of this architectural meshwork at the chromocenter, we generated stable cell lines expressing HMR and HP1a fused to an engineered ascorbate peroxidase from soybean (APEX2) under a copper inducible promoter to perform proximity biotinylation (Fig. 2A). Upon treatment of APEX2 expressing cells with biotin phenol and hydrogen peroxide a localized burst of diffusible biotin–phenoxyl radicals is generated. These radicals then react with nearby (< 20 nm) electron rich amino acid side chains leading to the biotinylation of neighboring proteins that can be subsequently purified and identified using shot gun mass spectrometry (26-32). We confirmed the expression of ectopic proteins by Western blotting using antibodies against HMR, HP1a and APEX2 (Fig. 2B). The APEX2 protein was fused to a double nuclear localization signal to determine the non-targeted nuclear proteome. To perform the biotinylation reaction at physiological protein levels, we used non-inducing conditions for HMR_AP_ expression but induced the expression of HP1a_AP_ (Fig. 2B and C). For the APEX2_NLS_ cell line we added copper sulfate to the medium before the biotinylation to induce the expression of APEX2_NLS_ and to match HP1a_AP_ expression levels. At the expression level used, HMR_AP_ localizes to centromeres, marked by the centromere-specific histone variant dCenpA, HP1a_AP_ occupies a domain in the nucleus, which coincides with endogenous HP1a staining and APEX_NLS_ localizes to the nucleus (Fig.2B), showing a proper nuclear localization of the fusion proteins under the conditions used.

**Figure 2:**
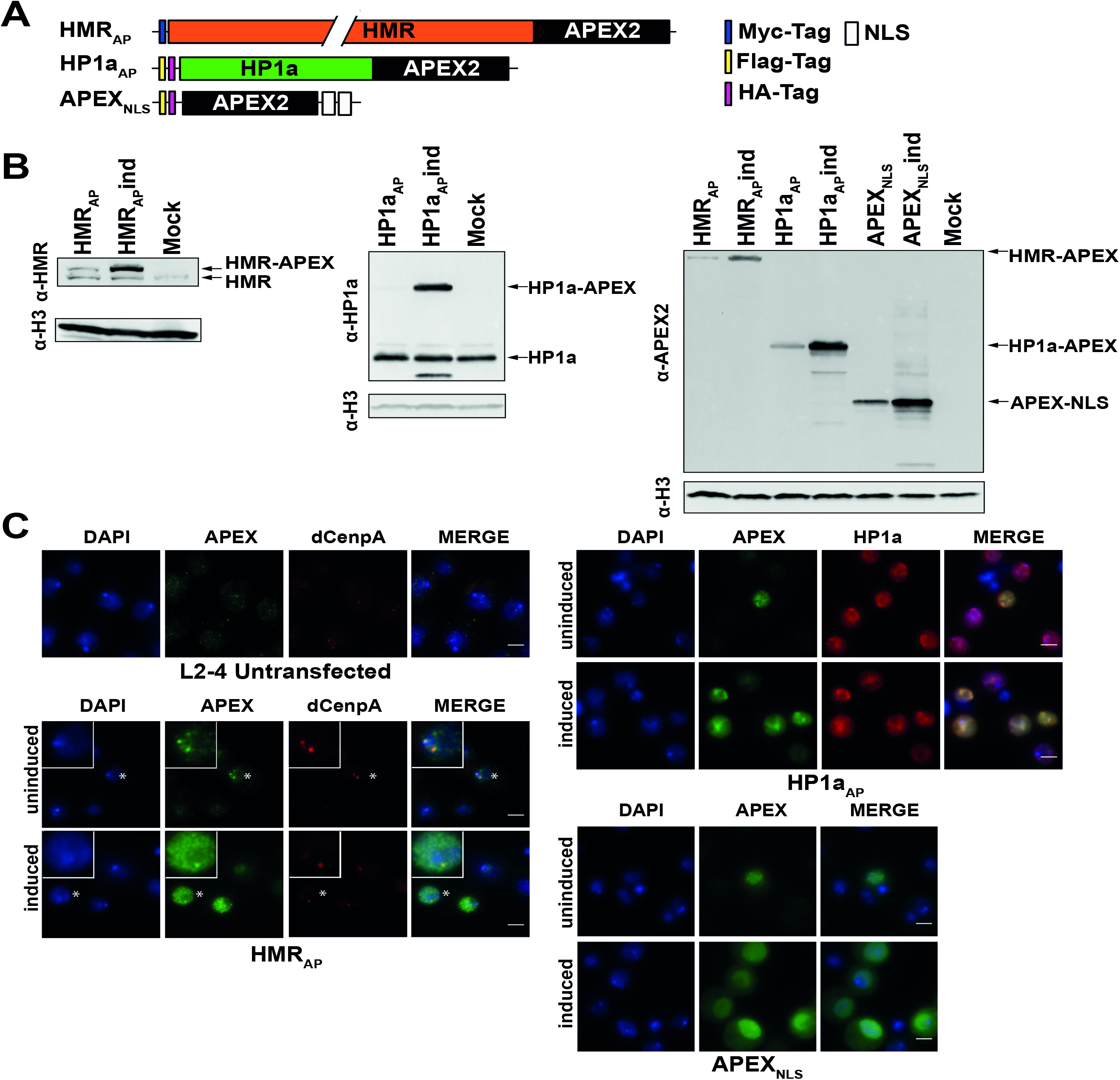
**(A)** APEX fusion constructs used for proximity based labeling of the chromocenter. **(B)** Western blots of whole cell lysates from stable cell lines, expressing HMR_AP_, HP1a_AP_ or APEX_NLS_. Antibodies used for Western blots were anti-HMR 2C10, anti-HP1a C1A9, anti-APEX2 20H10 and rabbit anti-H3 (for loading control). **(C)** Localization of APEX fusions in respective stable cell lines. Antibodies used are the same as for Western blotting in (**B**). For cells with centromere staining (indicated by an asterisk) the inlet shows an approx. 2.3-fold zoom of the nucleus. Shown are the stainings of cells with (induced) or without (uninduced) copper mediated induction of the expression of the corresponding fusion protein.

### HMR and HP1a APEX2 fusion proteins biotinylate defined nuclear domains

To analyze the localization of the biotinylation by the expressed fusion proteins, we stained the cell lines for APEX2 and biotin after performing an *in situ* biotinylation reaction (Fig. 3A). Consistent with the limited diffusion of the phenoxy radicals generated by APEX2, we observed a strong biotinylation signal colocalizing with the APEX2-fusion proteins (Fig. 3B). Cells that do not express nuclear APEX2 or cells that were not treated with biotin-phenol and hydrogen peroxide only showed background biotinylation signals. As the different cell lines expressed different levels of the APEX2-fusion protein and as HP1a has previously been shown to be very dynamic, we adjusted the concentration of biotin-phenol and the duration of hydrogen peroxide treatment to maintain proper localization while achieving a high degree of biotinylation of the neighboring proteome. While HMR_AP_ cells treated for 25 minutes with hydrogen peroxide at a high concentration of biotin-phenol (5 mM) showed a highly localized biotinylation (Supplemental Fig. S2), a similar treatment of HP1a_AP_ cells resulted in a broader biotinylation signal even though a reduced biotin-phenol concentration was used (0.5 mM) (Supplemental Figs. S2 and S3). This broader signal was not observed when HP1a_AP_ cells were treated for only 5 minutes with 0.5 mM biotin-phenol (Supplemental Fig. S3).

**Figure 3:**
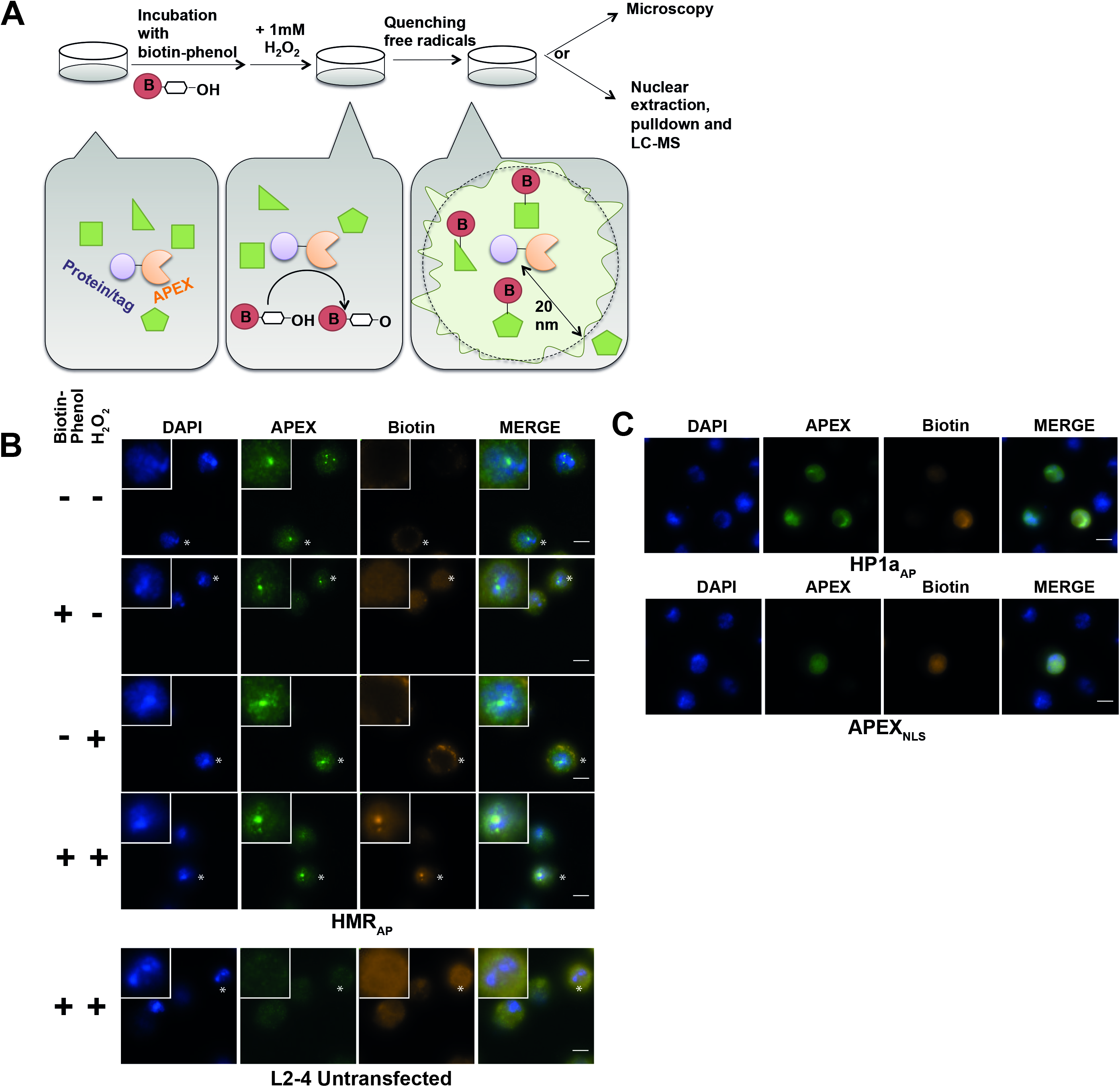
**(A)** Schematic display of a proximity based biotinylation experiment. **(B)** Biotinylation experiment performed for 1 minute with 5 mM biotin-phenol using the HMR_AP_ cell line (top four panels) in the absence of biotin phenol and hydrogen peroxide, the presence of either biotin phenol or hydrogen peroxide or both reagents. The bottom panel shows a treatment of non-APEX expressing L2-4 cells treated with biotin phenol and hydrogen peroxide. For selected cells (indicated by an asterisk) the inlet shows an approx. 2.3-fold zoom of the nucleus. **(C)** Proximity based biotinylation of the HP1a_AP_ and the APEX_NLS_ cell lines using 0.5 mM biotin-phenol. Stainings were performed with anti-APEX 20H10 antibody and anti-Streptavidin Alexa555. Scale bars represent 5 μm.

### Proteomic analysis of HMR and HP1a containing domains

Upon *in vivo* biotinylation, we compared the individual proteomes from cell lines expressing HMR_AP_, HP1_AP_ and APEX_NLS_ that were either treated with biotin-phenol and H_2_O_2_ or with DMSO only (Figure 4A). Using label free quantitation and a statistical analysis of the proximity-based proteome from at least 4 biological replicates we identified 406, 300 and 349 proteins that were specifically biotinylated in HMR_AP_, HP1a_AP_ and APEX_NLS_ expressing cells respectively (Supplementary Tab. S2). A comparative analysis of the isolated proximity proteomes revealed a strong overlap between the HP1a_AP_ and the APEX_NLS_ proteome (71%), which was not the case for HMR_AP_ and APEX_NLS_ (44%). The latter finding may reflect the lower expression level of HMR, the lower mobility and the more specific localization of HMR_AP_ compared to HP1a_AP_ (Fig. 3B). To further characterize the different proteomic composition within the three proximity proteomes we compared the enriched GO terms using the Gene Ontology Consortium tool (http://www.geneontology.org) (Supplementary Tab. S3). Consistent with its known function we found the GO terms such as chromatin silencing, histone modification and positive regulation of chromatin organization for HP1a_AP_. For HMR_AP_, we found the GO terms heterochromatin organization involved in chromatin silencing, telomere maintenance, mitotic sister chromatid segregation and nucleus organization, which were consistent with the observed phenotypes of HMR mutations in *Drosophila melanogaster* (13, 15, 22, 33, 34). This proximity-based proteome also strengthens the hypothesis that HMR localizes in between dCenpA and HP1a containing chromatin as components of both domains are found in proximity of HMR. Proteins that are biotinylated by all three factors were often highly abundant nuclear factors constituting the splicing machinery or structural proteins such as NLP, D1 or Lamin, which are likely to be in close proximity to most nuclear factors. As the long hydrogen peroxide treatment resulted in a lower HP1a_AP_ staining upon biotinylation, we also measured the HP1a_AP_ proximity proteome upon 5 minutes of biotinylation. Due to the shorter biotinylation time we identified less specifically biotinylated proteins (122). When comparing the proteins biotinylated after 5 minutes peroxide treatment with the ones isolated after 25 minutes of biotinylation we identified a higher percentage of previously characterized HP1a interactors after 25 minutes of biotinylation (35, 36) (Supplementary Fig. S4) suggesting that a longer biotinylation reaction covers a broader range of HP1a_AP_ containing domains. To identify proteins that specifically localize close to HMR_AP_ and HP1a_AP_ and are not distributed throughout the entire nucleus, we selected proteins that were preferentially biotinylated in the HMR_AP_ or the HP1a_AP_ cell line but not or only to a much lesser degree in the APEX_NLS_ line (Supplementary Tab. S2). As the HP1a_AP_ specific proximity proteome showed a large overlap with the proteome in proximity to APEX_NLS_, only very few chromatin associated factors were significantly closer to HP1a_AP_ than to APEX_NLS_ (Supplementary Table S1), which prevented us from further analyzing the HP1a_AP_ proximity proteome in great depth. For HMR_AP_, however we detected a substantial number of proteins being much closer to HMR_AP_ than to APEX_NLS_ and therefore displayed the proximity-based proteome of HMR_AP_ in the form of a network graph. In this graph individual proteins are displayed as nodes and previously published interactions as edges (Fig. 4C). Besides a large fraction of nucleolar proteins, we find the condensin and subunits of the cohesin complex, a large fraction of nuclear pore proteins, several insulator proteins, as well as a network of transcription factors including the previously identified hybrid incompatibility factor GFZF, as well as nucleosome remodeling factors and histone-modifying enzymes (Fig. 4C).

**Figure 4:**
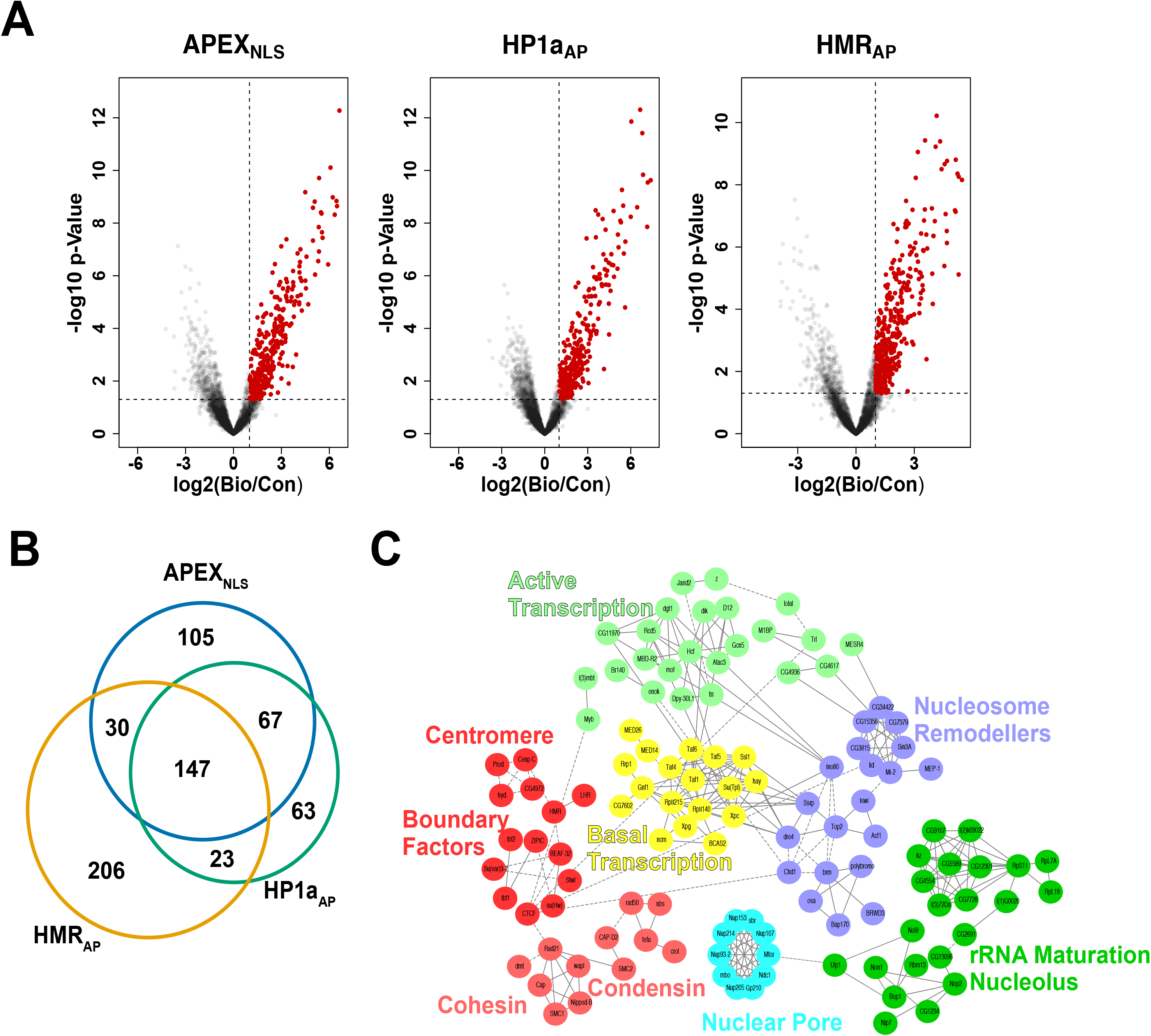
**(A)** Volcano plots of streptavidin pulldowns from extracts prepared from cells expressing APEX_NLS_, HP1a_AP_ and HMR_AP_ either mock treated or treated with biotin phenol and peroxide. Proteins specifically biotinylated in the treated samples are indicated in red. **(B)** Venn diagram of proteins found enriched in the individual purifications. **(C)** Network diagram of proteins specifically enriched in HMR_AP_ but not in APEX_NLS_. Solid lines were provided by the STRING databases using the highest stringency settings (0.9). Dotted lines were manually added based on information from Flybase.

### Cohesin and condensin factors reside in proximity to HMR

We were particularly intrigued by the proximity of HMR_AP_ to cohesin and condesin subunits as it has been recently shown that hybrid male flies that have higher levels of HMR show defects in chromosome condensation and integrity reminiscent of those observed in cells depleted of condensins or showing abnormal accumulation of cohesins (15). To validate the proximity of HMR with members of the cohesin complex, we compared the ChIP-sequencing profiles of HMR in S2 cells (20), the cohesin subunit Rad21 (vtd) and the condensin subunit CAP-H2 in Kc167 cells (37). Consistent with the proximity labelling result, approximately half (51%) of the HMR binding sites colocalize with binding sites of the cohesin complex and 20% with the CAP-H2 subunit of the condensin complex (Fig. 5A). The analysis of the genomic features of the HMR binding sites that overlap with Rad21 or CAP-H2 demonstrated that the HMR binding sites colocalizing with condensin or cohesin are more frequently found at promoter regions (Fig. 5B, left panel). By classifying chromatin domains using the five-state chromatin model (38) the HMR binding sites that colocalize with the condensin subunit CAP-H2 were more frequently found at active chromatin or HP1a containing heterochromatin (Fig. 5B, right panel).

**Figure 5:**
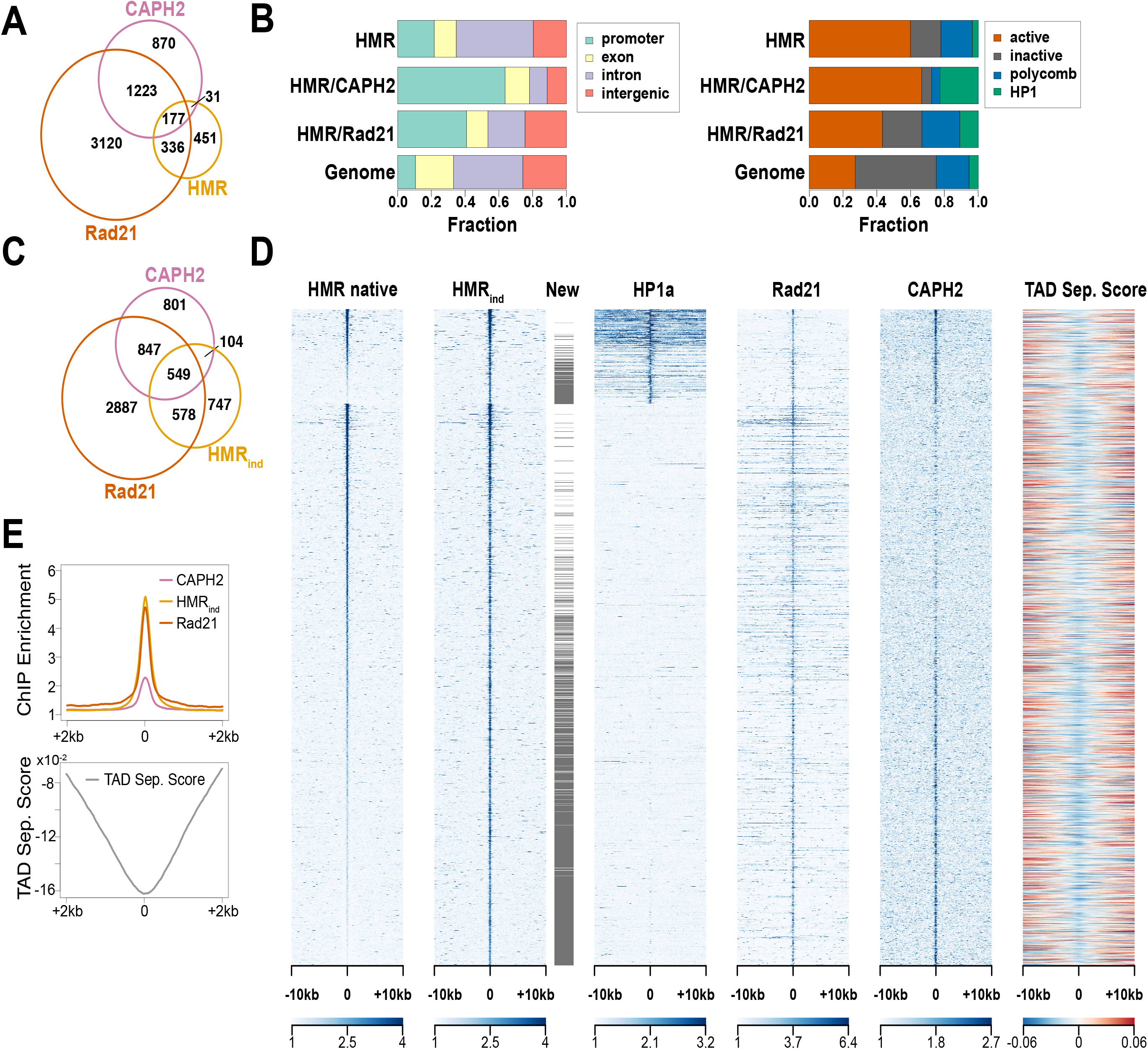
**(A)** Peak overlaps of HMR, Rad21 and CAP-H2. **(B)** Distribution of HMR, HMR/Rad21 and HMR/CAP-H2 peaks across different genomic regions and chromatin types. **(C)** Peak overlaps of overexpressed HMR, Rad21 and CAP-H2. **(D)** Heatmaps of HMR (20), overexpressed HMR, HP1a (20), Rad21, CAP-H2 ChIP-seqs (54) and TAD separation score (40). All the peaks are centered at HMR binding sites and clustering is performed as in (20). **(E)** Composite plot of overexpressed HMR, Rad21, CAP-H2 ChIP-seq signals and TAD separation score centered to the HMR peaks’ position. Peaks that are only detected when HMR is overexpressed are indicated as new.

To simulate the hybrid situation where increased levels of the complex containing HMR and LHR are found (13), we compared a HMR ChIP-seq analysis upon overexpression of HMR and LHR (39) with the condensin and cohesin profiles. Upon overexpression we detected 1314 additional HMR peaks, many of them colocalizing with binding sites of CAP-H2 (Fig. 5C). When we aligned ChIP-seq profiles along HMR binding sites (both native and induced), we found, that CAP-H2 ChIP enrichment is greater at HMR sites having high HP1a signals or at HMR sites appearing only upon overexpression (Fig. 5D). Rad21 ChIP enrichment in contrast is non-discriminating for classes of HMR binding. Architectural proteins, such as condensins and cohesins, were shown to cluster at TAD boundaries (37). We therefore wondered, whether HMR binding sites also reside there. Interestingly, HMR binding sites have local minima in TAD separation score (40) (Figs. 5D and E) suggesting that HMR localizes and potentially functions at genomic sites important for the 3D organization of chromatin.

## DISCUSSION

Interspecies hybrids are either sterile or lethal due to the incompatibility of their genomes (41, 42) or the incompatibility of the oocyte cytoplasm with a conflicting parental genome. Crosses between the closely related species *Drosophila melanogaster* and *Drosophila simulans* result in hybrid male lethality dependent on the presence of *D. mel* HMR and its interaction partner LHR from *D. sim*. More recently, gfzf, a gene encoding an essential FLWYTCH domain containing protein residing on the 3^rd^ chromosome of *D.sim*, has also been shown to contribute to male hybrid lethality (43). Though the effect of these factors on hybrid male lethality is well described, their molecular function is still largely unknown. HMR has been implicated in several cellular processes such as the silencing of transposable elements (13, 22, 48) or the faithful chromosome segregation and sister chromatid separation (13, 15, 33). HMR forms distinct foci that frequently overlap with the centromere in *D. mel* cell lines as well as in imaginal discs derived from *D.mel* larvae. An *Hmr* transgene carrying a mutation that affects its centromeric localization fails to kill male hybrids, suggesting that HMR’s mode of chromatin binding is causally related to the male lethality phenotype (13, 34). Our systematic microscopic analysis reveals that while most of the HMR foci colocalize with dCenpA this is not always the case. This suggests either a cell cycle regulated HMR-centromere interaction or a chromosome specific binding of HMR similar to the SATIII RNA, which has also been implicated in species formation (44) and only associates with chromosomes 2,3 and X (18). The overall localization of HMR may also be dependent on the cell type as we for example do not observe sizeable telomeric localization of HMR in S2 cells, while it is substantial in polytene chromosomes (13). Super resolution microscopy of the HMR containing nuclear domains furthermore revealed that HMR, dCenpA and dCenpC partially overlap within a nuclear domain that is clearly separated from HP1a containing heterochromatin. We interpret these findings as an indication of HMRs role in separating individual chromatin domains.

Within many organisms centromeres, pericentromeric heterochromatin and in some cases even telomeres cluster to form a nuclear domain or aggregate called chromocenter (24). These aggregates are frequently found at the nuclear periphery or around the nucleolus. Strikingly, we find HMR in proximity of nucleolar components and members of the nuclear pore suggesting that it might function in positioning the chromocenter within the nucleus after cell division.

Besides these structural components of the nucleus, the HMR proximity proteome also reveals its close localization to factors that separate functional chromatin domains. This is consistent with previous ChIP-seq mapping of HMR binding sites, which showed that a subset of HMR peaks border HP1a containing chromatin at promoters of active genes (20). Many of these HMR binding sites also colocalize with a condensin complex, two subunits of which we also find in close proximity to HMR. The proximity of HMR with condensin and cohesin at sites that separate different chromatin domains is consistent with the findings of Blum and colleagues (15) who described the *Hmr* phenotype as reminiscent to perturbations in cohesin or condensin function with frequent anaphase breakages at boundaries between eu- and heterochromatin. The striking discrepancy between the number of HMR binding sites observed in ChIP-seq experiments and the few histologically visible clusters suggest that HMR binding sites cluster *in vivo* at the chromocenter (Fig. 6). Such a role of clustered boundary regions in organizing nuclear 3D structure has also been observed in human as well as *Drosophila* cells (45, 46) suggesting a common mechanism for TAD organization. These domains not only play important function in the regulation of gene expression but also in reassembling the nuclear structure during the mitotic telophase. Increased levels of HMR, which is also observed in hybrids, results in an accumulation of HMR binding at previously unbound CAP-H2 binding sites which may in turn result in the formation of multiple novel clusters and a dysfunctional organization of the nucleus (47).

**Figure 6:**
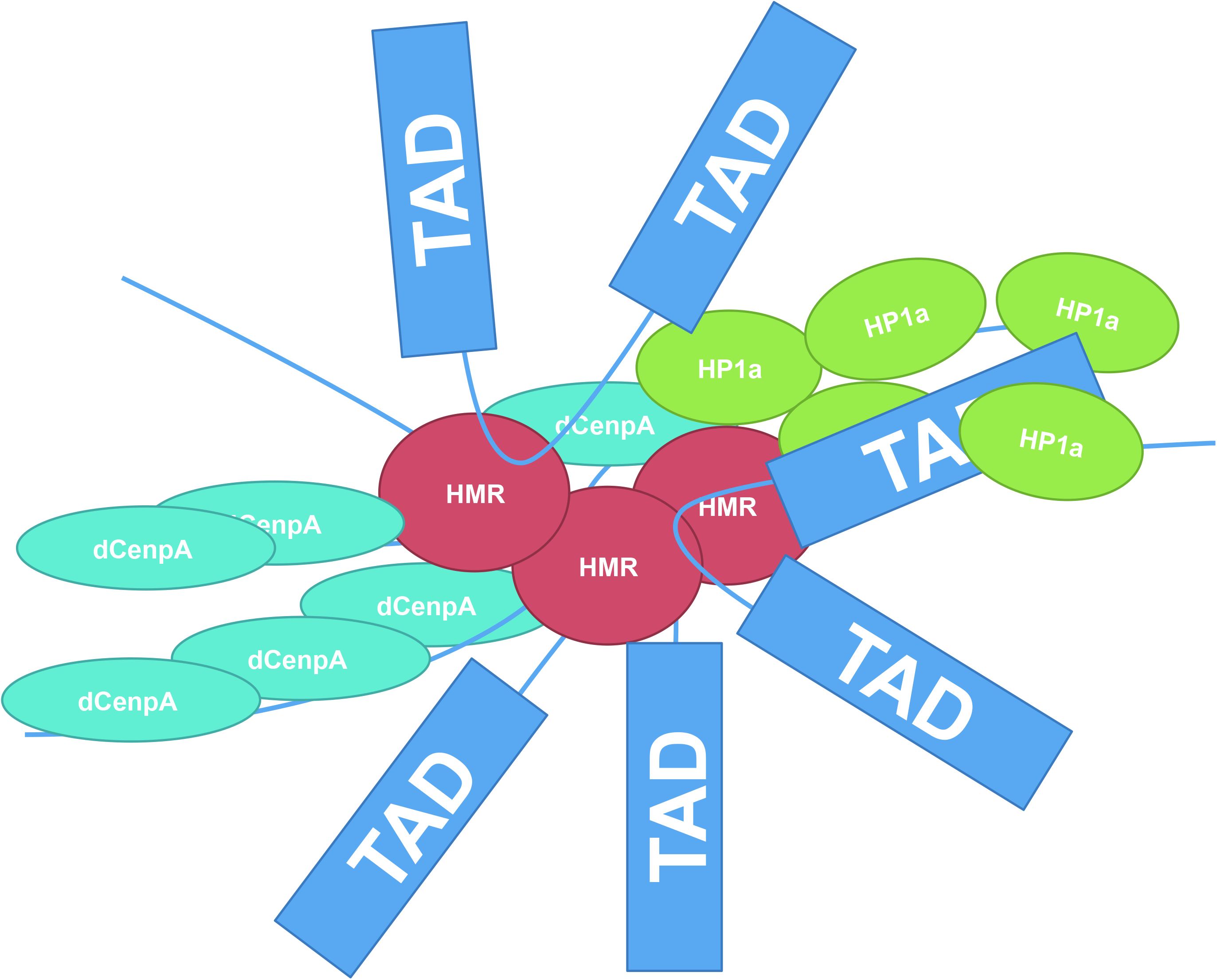
Model of organization of *Drosophila* interphase centromere. HMR and dCenpA protein domains are interdigitating, and HMR domains are often bordering dCenpA domains from HP1a domains. Some TAD boundaries, occupied by HMR, might be clustered at the border between centromeric and pericentric chromatins.

In summary, our results suggest that HMR is, in conjunction with a highly complex molecular machinery, involved in organizing the nuclear 3D structure. A failure of this tight regulation could then interfere with this well-orchestrated and self-organized assembly process leading to a catastrophic outcome such as hybrid lethality.

## MATERIAL AND METHODS

### Cloning

APEX fusions were cloned into pMT vector (13), which was cut with XbaI and NotI. GST-APEX was cloned into pGEX-6P-1 vector (13), cut with EcoRI and NotI. Cloning was performed with In-Fusion cloning kit (Clontech). The list of primers is available in the supplementary table S1.

### Cell culture, transfection and generation of stable cell lines

*Drosophila* L2-4 cells (13) were grown in Schneider medium supplemented with 10% fetal calf serum, penicillin and streptomycin at 26°C. To generate a stable cell line, 3-4 million of cells were transfected with 2 μg of plasmid mixed with XtremeGENE HP (Roche) transfection reagent according to manufacturer’s instructions. After transfection, cells were selected for 3 weeks with Hygromycin B (Invitrogen) at 100 μg/ml. Optional induction of cell lines with 250 μM CuSO_4_ was performed 12-24 hours before experiments.

### APEX proximity biotinylation, nuclear extraction and immunoprecipitation

For biotinylation cells were grown in roller bottles (Greiner) to density of 5*10^6^ cells/ml. For biotinylation in solution 5*10^8^-10^9^ cells were resuspended in 200 ml biotin-phenol/PBS or DMSO/PBS and incubated for 30 min at room temperature (RT). H_2_O_2_ was added to biotin-phenol treated cells to the concentration of 1 mM and cells were pelleted for 20 min at 250 g. The supernatant was aspirated, and cells were washed three times in quenching solution (10 mM sodium azide, 10 mM sodium ascorbate and 5mM Trolox in PBS) or (untreated cells) in PBS. The last washing step was performed in 15 ml falcons. Nuclei isolation and extraction was performed as in (48) with modifications. All buffers were supplemented with freshly added protease inhibitors. Cells were swelled in 3 packed cell volumes (PCV) (e.g. 2,1 ml) of hypotonic buffer (10 mM Tris pH 7.6, 10 mM NaCl, 1.5 mM MgCl_2_, 0.1 mM EDTA) on ice for 30 min. Cells were centrifuged 10 min 250 g, and the pellet was resuspended in 3 PCV of hypotonic buffer supplemented with 0,2% NP-40. Cells were rotated for 10 min at 4°C for the lysis of plasma membrane. Nuclei were pelleted at 1000 g for 10 min and washed with 2 ml of quenching solution supplemented with protease inhibitors. After the wash nuclei were centrifuged for 10 min with 1500 g and the nuclear pellet was snap-frozen in liquid nitrogen. Nuclei were resuspended in 3 ml Tris-Ex100 buffer (10 mM Tris pH 7.6, 100 mM NaCl, 1.5 mM MgCl_2,_ 0.5 mM EGTA and 10% v/v glycerol) + 1500 units Mnase + 1500 units Benzonase + 2mM CaCl_2_. Chromatin was digested for 20 min at 26°C and the reaction was stopped by addition of EDTA and EGTA to 10 mM on ice. Nuclei were disrupted by Dounce homogenization using 10 strokes with a tight-fitting pestle. For chromatin extraction and solubilization, NaCl (to 600 mM) and detergents (Triton X-100 to 1%, sodiumdeoxycholate (SOD) to 0.5% and SDS to 0.1%) were added and chromatin was incubated for 1h at 4°C. Nuclear extracts were cleared by centrifugation 20 min 10000 g and dialyzed for 4 hours at 4°C through 3.5 MWCO Millipore membranes against Tris-Ex100 with detergents and without glycerol, supplemented with 0.2 mM PMSF and 1 mM DTT. The obtained nuclear extract was snap-frozen. For streptavidin purification, nuclear extract with freshly added protease inhibitors was mixed with 500 μl of Pierce streptavidin magnetic beads, prewashed two times in Tris-Ex100 buffer with detergents. Immunoprecipitation was performed for 1,5 hours at RT. Beads were washed two times with Tris-Ex100 buffer with detergents (with protease inhibitors), once with 2 M Urea 10 mM Tris, and again twice with Tris-Ex100 with detergents and protease inhibitors.

For biotinylation on plates, 800 ml of 5*10^9^ million/ml cells were adhered on 40 15-cm plates for mammalian cells for 1 hour. Cells were incubated in biotin-phenol/PBS (20 plates) or DMSO/PBS (20 plates) for 30 min, and H_2_O_2_ was added to biotin-phenol treated cells to the concentration of 1 mM for 5 min. Solution was aspirated and quenching solution (or PBS for untreated cells) was added. Cells were scraped off, washed one more time in quenching solution and subjected to nuclear extraction as described above.

### Mass spectrometry

Beads from APEX pulldowns were washed 3 times with 500 μl of 50 mM Tris pH 7.5, 4M urea, and on-bead digestion was performed. Reduction was completed in 500 μl of 20 mM DTT in 50 mM Tris, 2M urea pH 7.5 with Lys C 450 ng/sample at 27°C for 1h. Alkylation followed with 50 mM iodoacetamide for 1 h 25°C, shaking 900 rpm, and was stopped by addition of DTT to 10 mM final concentration. The samples were incubated at 25°C shaking 900 rpm for 2 more hours. 300 μl water was added to reduce urea concentration to 1.5 M, and 1.5 μg trypsin and 2 mM final concentration CaCl_2_ were added for overnight incubation shaking 900 rpm. In the morning another 1.5 μg of trypsin was added, and the sample was incubated for another 4 hours while shaking. After collection of the supernatant, the beads were washed 2 times with 100 μl of 20 mM Tris 50 mM NaCl 25% ACN in order to elute loosely-bound tryptic peptides from the beads. Washes were combined with supernatant, evaporated at less than 28°C, and desalted. The second elution of remaining peptides from the beads was performed with 300 μl 0.05% SDS, 0.1% formic acid (FA) at 80°C for 10 min. The second elution was evaporated and subjected to HILIC chromatography. After desalting and HILIC both elutions were combined, evaporated and resuspended in 45 μl 0.1% FA. Peptide mixtures were subjected to nanoRP-LC-MS/MS analysis on an Ultimate 3000 nano chromatography system coupled to a QExactive HF mass spectrometer (both Thermo Fisher Scientific) in 2-4 technical replicates (5 μl each). The samples were directly injected in 0.1% FA onto the separating column (120 x 0.075 mm, in house packed with ReprosilAQ-C18, Dr. Maisch GmbH, 2.4 μm) at a flow rate of 0.3 μl/min. The peptides were separated by a linear gradient from 3% ACN to 40% ACN in 50 min. The outlet of the column served as electrospray ionization emitter to transfer the peptide ions directly into the mass spectrometer. The QExactive was operated in a TOP10 duty cycle, detecting intact peptide ion in positive ion mode in the initial survey scan at 60,000 resolution and selecting up to 10 precursors per cycle for individual fragmentation analysis. Therefore, precursor ions with charge state between 2+ and 5+ were isolated in a 2 Da window and subjected to higher-energy collisional fragmentation in the HCD-Trap. After MS/MS acquisition precursors were excluded from MS/MS analysis for 20 seconds to reduce data redundancy. Siloxane signals were used for internal calibration of mass spectra.

### Protein MaxQuant search

The raw data files of APEX pulldowns were analysed with MaxQuant version 1.5.3.12 against dmel-all-translation-r6.08.fasta database from Flybase. All the parameters were set to default except choosing “Match between runs” and LFQ and iBAQ quantitations. Technical replicates were assigned to one experiment (biological replicate).

### Data sources

ChIP-seq datasets were available from GEO with the following accession numbers: GSE86106 (HMR native and HP1a), GSE118291 (HMR induced), GSE54529 (Rad21 and CAPH2). 5-state chromatin domains were taken from (43) from which “yellow” and “red” were merged to active domains. Genomic regions (promoters, exons etc.) are based on the Flybase annotation (version r6.17). TAD separation scores derived from http://chorogenome.ie-freiburg.mpg.de/data_sources.html. Genomic coordinates from the dm3 release were converted to dm6 using the liftover tool from UCSC.

### ChIP-seq analysis

Raw reads were aligned to the reference genome (UCSC dm6) using bowtie2 (version 2.2.9) and quality filtered by samtools (version 1.3.1). ChIP-seq profiles were created by Homer (version 4.9) and normalized by the number of reads and the corresponding input. Peaks were called by Homer with parameters -style factor -F 2 -size 200 (except for Rad21 -F 4). Peak overlaps were plotted as Venn diagrams using the Vennerable R package (version 3.1.0.9). HMR peaks were sub-grouped by the overlap with CAPH2 or Rad21 peaks. Such peak groups were characterized by genomic features (promoters, exons, etc.) and chromatin domains (active, etc.) by taking the size of the intersecting genomic ranges between the peaks and each feature and dividing by the sum of the intersecting range sizes. ChIP-seq profiles were aligned to the pool of native and induced HMR peaks and visualized as heatmaps or as average composite plots. Heatmaps were ordered by HMR native enrichment and grouped by HP1a enrichment. Figures were created by R base graphics.

### Mass spectrometry analysis

Log2-transformed LFQ values were imputed with the impute.MinProb function (imputeLCMD R package v2.0) with parameter q = 0.05 and normalized by median normalization. Moderated t-statistics were computed by fitting a linear model with empirical Bayes moderation using the limma R package (version 3.34.9). The data were visualized as volcano plots using R graphics. Code is available upon request. GO-term analysis was done using Gene Ontology Consortium (http://geneontology.org) tool excluding redundant GO terms. The protein-protein interaction network was visualized using Cytoscape. Protein-protein interaction data were taken from the STRING database (49) using experiments, databases, gene fusion and co-occurrence as interaction sources and minimum required interaction score of 0.9. Additional interactions were taken from the database Flybase and indicated as dotted lines.

### Immunofluorescent staining

Cells were adhered on coverslips for 30 min, washed with PBS and fixed in 0.3% Triton X-100/3.7% formaldehyde/PBS for 12 min at RT. After wash with PBS cells were permeabilized in 0.25%Triton/PBS on ice for 6 min. Cells were rinsed 2 times and washed again twice with PBS, following blocking in Image-iT FX signal enhancer (Invitrogen) for 45 min at RT. Primary antibodies, diluted in 5% normal goat serum (NGS) (Dianova), were incubated with coverslips 1 h at RT (or at 4°C overnight for high resolution microscopy). Coverslips were washed with 0.1% Triton X-100/PBS and PBS and incubated with secondary antibody in 5% NGS 1 h at RT. Following a wash in 0.1% Triton X-100/PBS and 2 washes in PBS, cells were stained in DAPI/PBS (200 ng/mL or 50 ng/mL for high resolution microscopy) and washed again in PBS. Cells were mounted in VECTASHIELD (Vector Labs) or ProLong™ Diamond Antifade (Thermo Fisher Scientific), for epifluorescence and confocal or super resolution microscopy, respectively.

For biotinylation followed by immunofluorescence on coverslips, 10^6^ cells was adhered on coverslips for 30 min, incubated with biotin-phenol/PBS or DMSO/PBS for 30 min, followed by (optional) addition of 1mM H_2_O_2_ for denoted time. Cells were next washed in quenching solution and subjected to immunofluorescent staining as described above. Biotinylation in solution was performed with 10^6^ cells in 200 μl biotin-phenol/PBS or DMSO/PBS and 1 mM H_2_O_2_, added for denoted time. Cells were pelleted during biotinylation 250 g, resuspended in 200 μl quenching solution, adhered on coverslips for 15 min and processed for IF as described above.

Image acquisition was performed on a Zeiss Axiovert 200 epifluorescence microscope with a CCD Camera (AxioCam MR, Zeiss). Confocal microscopy was performed at the core facility bioimaging of the Biomedical Center with an inverted Leica SP8X WLL microscope, equipped with 405 nm laser, WLL2 laser (470 - 670 nm) and acousto-optical beam splitter. Gated-STED Images were acquired with a 100×1.4 objective, image pixel size was 24-25 nm. The following fluorescence settings were used: DAPI (excitation 405 nm; emission 415-470 nm), Alexa Fluor 594 (590 nm; 600-625) and Abberior STAR 635P (635; 645-720). Recording was done line sequentially to avoid bleed-through and channel misalignment because of drift. Signals were recorded with hybrid photo detectors (HyDs) in counting mode. STED and confocal images were deconvolved using Huygens 17.10 p2. All images were processed using ImageJ.

### Western blotting of whole cell extracts

15 million cells from each cell line were collected, washed two times in PBS and resuspended on ice in 80 μl RIPA buffer with freshly added protease inhibitors and 30 units benzonase. Lysates were left on ice for 30 min, and afterwards 20 μl of 5x Laemmli buffer was added. Lysates were boiled 10 min 96°C before loading on the gel (10 μl).

### Antibodies

For immunofluorescence and western blotting, rat anti-HMR 2C10 antibody (Helmholtz Zentrum München, (13)) was used in dilution 1:25 (or 1:5 for high resolution microscopy); rat anti-dCenpA 7A2 (Helmholtz Zentrum München, (8)) 1:100 (or 1:50 for high resolution microscopy); rabbit anti-dCenpA (Immunofluorescence grade, Actif Motif) 1:500 (or 1:250 for high resolution microscopy); mouse anti-HP1a C1A9 (50) 1:100; anti-APEX2 20H10 (raised in this study, Helmholtz Zentrum München) 1:50; Streptavidin-Alexa555 (Thermo Fisher Scientific) 1:400; rabbit anti-histone H3 (Abcam) 1:3000, mouse anti-FLAG (Sigma M2, 1mg/ml) 1:100. Rabbit anti-dCenpC antibody was kindly provided by Christian Lehner and used in dilution 1:5000 (1:1000 for high resolution microscopy).

### Antibody generation

Wistar rats were immunized subcutaneously (s.c.) and intraperitonially (i.p.) with 50 μg of GST-APEX fusion protein dissolved in 500μl PBS, 5 nmol CpG2006 (TIB MOLBIOL) an equal volume of incomplete Freund’s adjuvant. 6 weeks after immunization a 50 μg boost injection was applied i.p. and s.c. three days before fusion. Fusion of the splenic B cells and the myeloma cell line P3X63Ag8.653 was performed using polyethylene glycol 1500 according to standard protocols (51). Hybridoma supernatants were tested by solid-phase enzyme-linked immunoassay (ELISA) using the recombinant GST-fusion protein and verified by Western blotting of whole cell extracts from APEX2 fusions-expressing cell lines (Fig. 2B). Hybridoma cell line from specifically reacting supernatants were cloned twice by limiting dilution. Experiments in this study were performed with clone 20H10 (rat IgG2a/κ).

## DATA AVAILABILITY

The mass spectrometry proteomics data have been deposited to the ProteomeXchange Consortium via the PRIDE (52) partner repository with the dataset identifier PXD011310.

## ACKNOWLEDGEMENT

We thank the Christian Lehner and Sarah Elgin for the Rabbit anti dCenpC and the mouse anti-HP1a antibody respectively. We furthermore would like to thank Nitin Phadnis and Patrick Heun for critical comments on the manuscript as well as as Irene Vetter for cloning the HP1a-APEX2 fusion construct, Ignasi Forne for mass-spectrometry analysis, Tobias Straub for initial bioinformatic analysis and the entire Imhof group/Becker department for helpful discussions.

## FUNDING

This work was supported by the grants from the Deutsche Forschungsgemeinschaft (DFG) to AI (CIPSM and AI23/9-1), NK and AL (Graduate School of Quantitative Biosciences Munich (QBM)). Funding for open access charges: DFG CRC 1064/Z03

## CONFLICT OF INTEREST

The authors state no conflict of interest

## SUPPLEMENTARY DATA

**Figure S1:** Confocal microscopy images of L2-4 cells stained with rabbit anti-dCenpA, anti-HMR 2C10 and anti-HP1a C1A9 antibodies. The highest intensities graph was built in ImageJ and normalized to one of the maximum peaks.

**Figure S2:** Biotinylation experiment performed for 25 minutes using the HMR_AP_, HP1a_AP_ and APEX_NLS_ cell lines. Controls are shown in the right panel. Stainings were performed with anti-APEX 20H10 antibody and anti-Streptavidin Alexa555. For selected cells (indicated by an asterisk) the inlet shows an approx. 2.3x zoom of the nucleus. Scale bars represent 5 μm. Exposure for APEX and biotin is indicated in white.

**Figure S3:** Time-course biotinylation experiment performed for the HP1a_AP_ cell line. Stainings were performed with anti-APEX 20H10 antibody and anti-Streptavidin Alexa555. Scale bars represent 5 μm.

**Figure S4:** Volcano plots of streptavidin pulldowns from extracts of cells expressing HP1a_AP_ either mock treated or treated with biotin phenol and peroxide (for 5 and 25 minutes). Previously reported HP1a interactors, enriched in the pulldowns, are highlighted in green.

**Table S1:** List of primers used for cloning of Myc-HMR-APEX, Flag-HA-HP1a-APEX, Flag-HA-APEX-NLS and GST-APEX constructs.

**Table S2:** Lists of proteins, enriched in HMR_AP_, HP1a_AP_ and APEX_NLS_ pulldowns from treated against untreated cells, as well as proteins enriched in HMR_AP_ and HP1a_AP_ pulldowns against APEX_NLS_.

**Table S3:** GO term analysis of proteins enriched in HMR_AP_, HP1a_AP_ and APEX_NLS_ pulldowns from treated against untreated cells. GO terms found in HMR_AP_ and HP1a_AP_ but not APEX_NLS_ are highlighted in light green.

**Movie S1:** 3D reconstruction from confocal HP1a (green) and super resolution STED HMR (red) and dCenpA (blue) microscopy.

